# The effects of pressure on anticipatory postural adjustments during a jump shot in basketball

**DOI:** 10.1101/2024.05.10.593492

**Authors:** Kiyohiro Konno, Atsushi Itaya, Tomohiro Kizuka, Seiji Ono

**Author notes:** **Declarations of interest:** None.

## Abstract

**Background:** Anticipatory postural adjustments (APAs) play an important role in feedforward control of dynamic movement, such as jump shot performance in basketball. It is known that jump shot performance declines under pressure from a defender (shot blocker).

**Research question:** Does pressure from a shot blocker affect APAs of a jump shot in basketball? Is jump shot performance in basketball associated with APAs?

**Methods:** Fourteen healthy male university basketball players performed jump shots under pressure and non-pressure (free) conditions by a shot blocker. Using a force plate, the APAs were defined as occurring until the point of thrust (TH) phase, and ground reaction force (GRF) and center of pressure (COP) at that moment were assessed. To assess jump shot performance, the maximum GRF during the TH phase (TH_max_), jumping height, and success score of the shot (accuracy score: AS) were measured by using the vertical component of the force plate. Two-way ANOVA examined the effects of the phase and condition on APA duration. Pairwise t-tests analyzed pressure effects on subjective pressure intensity, AS, and kinetics measures. Relationships between condition changes in AS and COP or GRF variables were assessed via Pearson correlations. Differences between the pressure and free conditions were denoted by Δ.

**Results:** The APA duration was shorter under the pressure condition compared to the free condition. The TH_max_ and jump height values were greater under the pressure condition relative to the free condition. Across conditions, changes in COP variables were significantly and negatively correlated with AS.

**Significance:** The results of this study suggest that pressure from a shot blocker shortens APA duration and decreases jump shot performance. Additionally, by measuring players’ APAs and evaluating relationships with jump shooting proficiency, APA variables could potentially serve as indicators of skill expertise.

## 1. Introduction

Anticipatory postural adjustments (APAs) are an essential component of postural control [1]. The central nerves system (CNS) generates APAs by predicting perturbations from voluntary movements [2,3] to minimize effects on posture and enhance stability [4,5]. Hence, APAs, as a feedforward control, are essential for stable postural regulation during voluntary movements. Several previous studies have shown that APAs are associated with upward jump motions. For example, Pellec and Maton (1999) have revealed that APAs are backward shifts of the center of pressure (COP) during the unloading phase (UL phase) of the jump movements[6]. Another study has indicated that the CNS predicts perturbations not only from voluntary movement onset but also from overall sequences of motions [7]. APAs have been reported to play several roles in voluntary movements in previous studies. The jump shot motion that incorporates a countermovement jump utilizing stretch-shortening cycles may contribute to upward propulsion and increased jump height [8]. Additionally, several previous studies have demonstrated that postural balance is associated with shooting accuracy [9–11]. In other words, APA in postural control would play an important role in performance in various kinds of sports.

In basketball, shooting skills are extremely important and directly impact team winning [12,13]. Among various shot techniques, jump shots are regarded as the most efficient [13]. The shot motion in basketball involves preparation phase [14]. Especially in jump shots involving a jump, postural control during the preparation phase substantially influences jump shot performance. During basketball games, offensive players often execute jump shots while guarded by opponents trying to block shots [15]. Shot success rates decline when shooting against defenders attempting blocks compared to open shots [16,17]. Moreover, previous studies have revealed that defensive pressure induces quicker shooting motions and longer air times [15,17]. However, several issues regarding feedforward postural control in jump shot performance against defenders remain unclear.

Previous studies have revealed shorter APA duration under simple reaction time compared to self-paced instructions [18,19]. However, it remains unclear whether APAs change under pressure from the shot blocker. The purpose of this study is to examine whether APAs change under pressure from the shot blocker. Secondly, this study also examines how the APAs under pressure from the shot blocker influence the jump shot performance.

## 2. Materials and Methods

### 2.1. Participants

Fourteen healthy male university basketball players (age: 20.54 ± 1.22 years, height: 175.15 ± 7.80 cm, body weight: 66.62 ± 8.67 kg, years of experience: 10.77 ± 2.49 years) participated in this study. Prior to participation, the participants completed a questionnaire regarding age, height, body weight, years of basketball experience, injury history, and dominant leg. Individuals with a history of hip, knee or ankle injury within the past 6 months were excluded. Based on the questionnaire, all participants were right-leg dominant.

Prior to the experiment, the procedures of this study were explained to the participants both verbally and in writing in accordance with the Declaration of Helsinki. Written informed consent was obtained from all participants.

### 2.2. Experimental protocol

The participants performed under pressure (with a rushing shot blocker) and free (without a shot blocker) conditions to examine postural adjustments under pressure from a shot blocker. In this study, the shot blocker stood 3 m in front of the participant. the examiner stood 3 m to the left of the participant. In the pressure condition, when the examiner passed the ball to the participant, the shot blocker rushed to block the participant’s shot.

Before performing the task, each participant completed a 5-min warm-up. The participants stood on a force plate placed at the free throw line and maintained a shooting stance for 2 s, followed by 24 randomized trials (12 trials per condition) of the free and pressure conditions. To avoid fatigue and adaptation to the pressure from the shot blocker, the participants were instructed to sit in a chair placed 1 m behind the free throw line after each trial. During the break, a visual analog scale (VAS) was used to evaluate the degree of pressure perceived by the participants in each trial.

In the free condition, after maintaining a stationary stance on the force plate, the participants were instructed to receive a pass from the examiner 3 m to the left and immediately perform a jump shot. In the pressure condition, the procedure was the same as the free condition until receiving the pass. When the examiner passed the ball, the shot blocker in front of the participant rushed to block the shot. The participants were instructed to perform a jump shot while under pressure from the shot blocker.

### 2.3. Data acquisition and processing

To determine the starting point of each trial, an LED for synchronization (FMT-LED4, 4 Assist), 5-channel time counter (4 Assist), and a PC application (TimeCounter, Version 1.10n, 4 Assist) were used. The ground reaction force (GRF) and center of pressure (COP) during the free and under pressure conditions were measured using a force plate (500 × 600 mm, 9260AA6, Kistler). Signals from the force plate were sent through an A/D converter (G-FORCE, 4 Assist) and recorded on a PC at a sampling frequency of 1000 Hz using data collection software (A-Cap version 1.0.2, 4 Assist).

The signals from the force plate were smoothed using a 4th order low-pass Butterworth digital filter with a cut-off frequency of 10 Hz. The vertical component was used for analysis. Lift-off during the jump shot (<30 N) was identified from the collected GRF waveform. The 1s period with the smallest standard deviation from the start of data collection to lift-off was identified, and the average value during this period was defined as the participant’s body weight. The GRF data were normalized by the body weight. Similarly, the mean values in the medial-lateral and anterior-posterior directions were calculated from the 1s COP waveform. Using these mean values as the reference coordinates, the coordinate system of the COP waveform from the start of the trial to lift-off was determined.

Signal processing from the force plate and calculation of analysis parameters were performed using original software written in Scilab 6.1.1 (ESI Group, CeCILL) script language.

### 2.4. Data analysis

#### 2.4.1. Subjective pressure intensity

The visual analog mood scale (VAS) was used to assess the degree of pressure perceived by the participants in each trial. The VAS is able to measure emotional states in a quick, reliable and relatively sensitive manner [20]. The VAS consisted of a 100 mm horizontal line ranging from 0 (no pressure at all, left end) to 100 (extremely high pressure, right end). VAS data were quantified by measuring the distance from the left end to the vertical line (mm) marked by the participant.

#### 2.4.2. Accuracy score (AS) of jump shot

To measure the effect of pressure more accurately from the shot blocker on shooting accuracy, the following scoring system was used: 3 points if the basketball entered the net without touching the rim or backboard, 2 points if the ball entered after touching the rim or backboard, and 1 point if the ball missed after touching or not touching the rim or backboard [21].

#### 2.4.3. Kinetics data

For the GRF, the jumping motion during the jump shot was divided into three phases. Going back in time from take-off, the period when the GRF before take-off exceeded body weight (>1) was defined as the thrust phase (TH phase). Before the TH phase, there was a period with less than body weight loading (Fig. 1A). This period was defined as the unloading phase (UL phase). Further, a slight loading was observed just before the UL phase, which was defined as the pre-loading phase (PL phase). In this study, the preceding UL and PL phases during takeoff preparation were defined as APAs. The duration of each phase was calculated as PL duration, UL duration and TH duration. The maximum GRF during the PL phase, minimum GRF during the UL phase, and maximum GRF during the TH phase were measured as PL_max_, UL_min_ and TH_max_, respectively (Fig. 1A). Based on the aerial time (<30 N period) t [s], the jump height of the jump shot was calculated as:

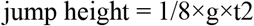

The g in the above formula is gravitational acceleration, using an approximate value of 9.81. For the COP, the COP was analyzed according to the phases determined by the GRF. The COP excursion from the PL phase until the end of the UL phase was calculated (APA excursion). This was divided by the time taken to obtain the APA sway velocity. Similarly, the sway ranges in the medial-lateral and anterior-posterior directions during this period were calculated as the APA medial-lateral range (APA ML range) and APA anterior-posterior range (APA AP range), respectively (Fig. 1B).

**Figure 1.**
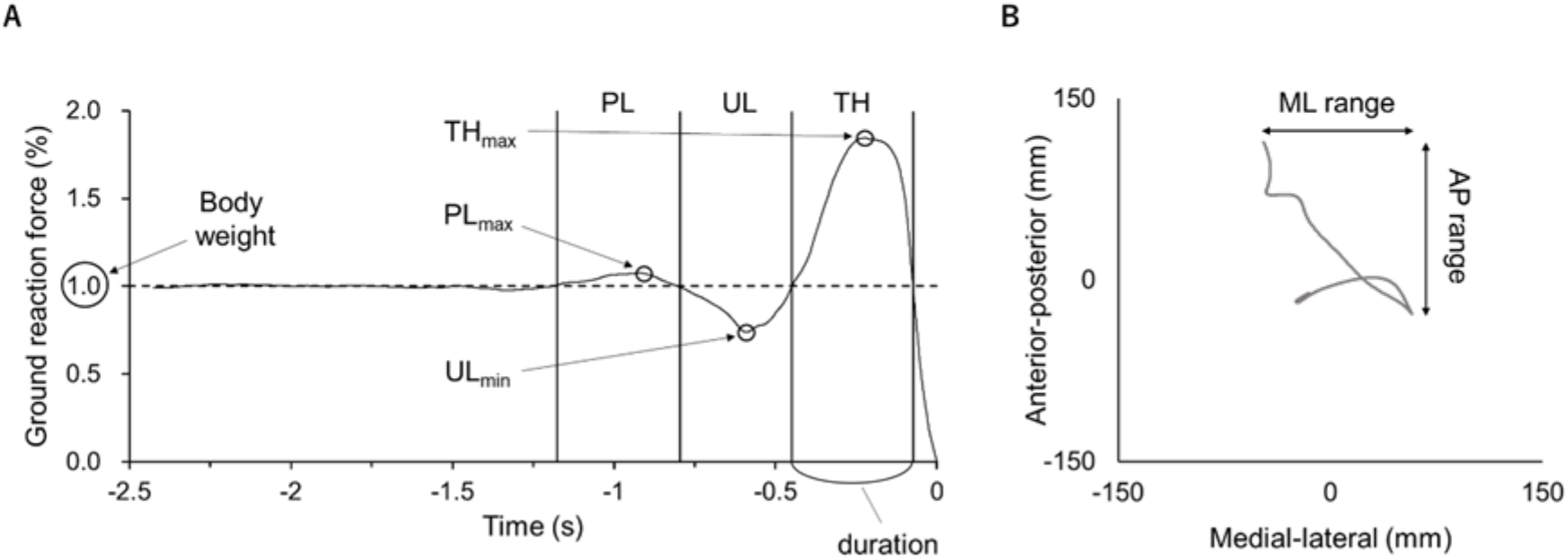
Representative individual data in the jump shot movement shows GRF and COP. (A) the GRF waveform of the participant’s jump shot up to take-off. The GRF data were normalized by the body weight. Each phase was defined as follows: the pre-loading (PL) phase is before the unloading (UL) phase with slight loading, the UL phase is the period below body weight, and the thrust (TH) phase is from take-off until GRF exceeds body weight. The duration of each phase was calculated as PL_dur_, UL_dur_ and TH_dur_. The maximum GRF during the PL phase, minimum GRF during the UL phase, and maximum GRF during the TH phase were measured as PL_max_, UL_min,_ and TH_max_, respectively. (B) the COP excursion in the jump shot up to take-off. The ranges from minimum to maximum values on the medial-lateral and anterior-posterior axes of the COP excursion were defined as the ML range and AP range, respectively.

#### 2.5. Statistical analysis

All results are reported as mean ± standard deviation. Statistical analysis was performed in jamovi (The jamovi project, 2022, ADPL 3).

To examine the effect of pressure on APA duration, the two-way analysis of variance ANOVA with factors of phase (PL and UL) and condition (free and pressure) was conducted. To investigate the effect of pressure from the shot blocker on each parameter (Subjective pressure intensity, AS and Kinetics data), the pairwise t test was conducted between the free and pressure conditions. To examine the relationship between changes (f-p) in AS and changes (f-p) in GRF and COP variables between conditions, the Pearson product-moment correlation coefficient was calculated. Differences between the pressure condition and free condition were denoted with Δ. Statistical significance was set at p < .050, and only those results that were significant were reported here.

## 3. Results

### 3.1. Subjective pressure intensity

The VAS data after each trial showed that the level of pressure intensity was significantly higher in the pressure condition than in the free condition for the participants in this study (t(13)= 20.497, p < .001).

### 3.2. AS of jump shot

When participants conducted the jump shot in the pressure condition their AS was lower compared to when they conducted the jump shot in the free condition. The average AS was 18.214 for the pressure condition and 22.571 for the free condition. The pairwise t-test revealed that this difference was significant (t(13) = 3.3004, p = .006).

### 3.3. Kinetics variables

GRF and COP of participants are shown in Table 2. For the APA phase, the repeated-measures two-way ANOVA with factors of phase (PL, UL) and condition (free, pressure) was conducted about APA duration (Fig. 2A). There was a significant main effect of phase on APA duration (F1,13 = 13.224, p = .003), as well as a significant main effect of condition on APA duration (F1,13 = 8.036, p = .014). No interaction between phase and condition was found in APA duration. The duration in the UL phase (free: 0.283±0.0256, pressure: 0.248±0.0220) was longer than that in the PL phase (free: 0.169±0.0199, pressure: 0.148±0.0223), and the duration in the pressure condition (PL: 0.148±0.0223, UL: 0.248±0.0220) was shorter than that in the free condition (PL: 0.169±0.0199, UL: 0.283±0.0256).

**Table 1.**
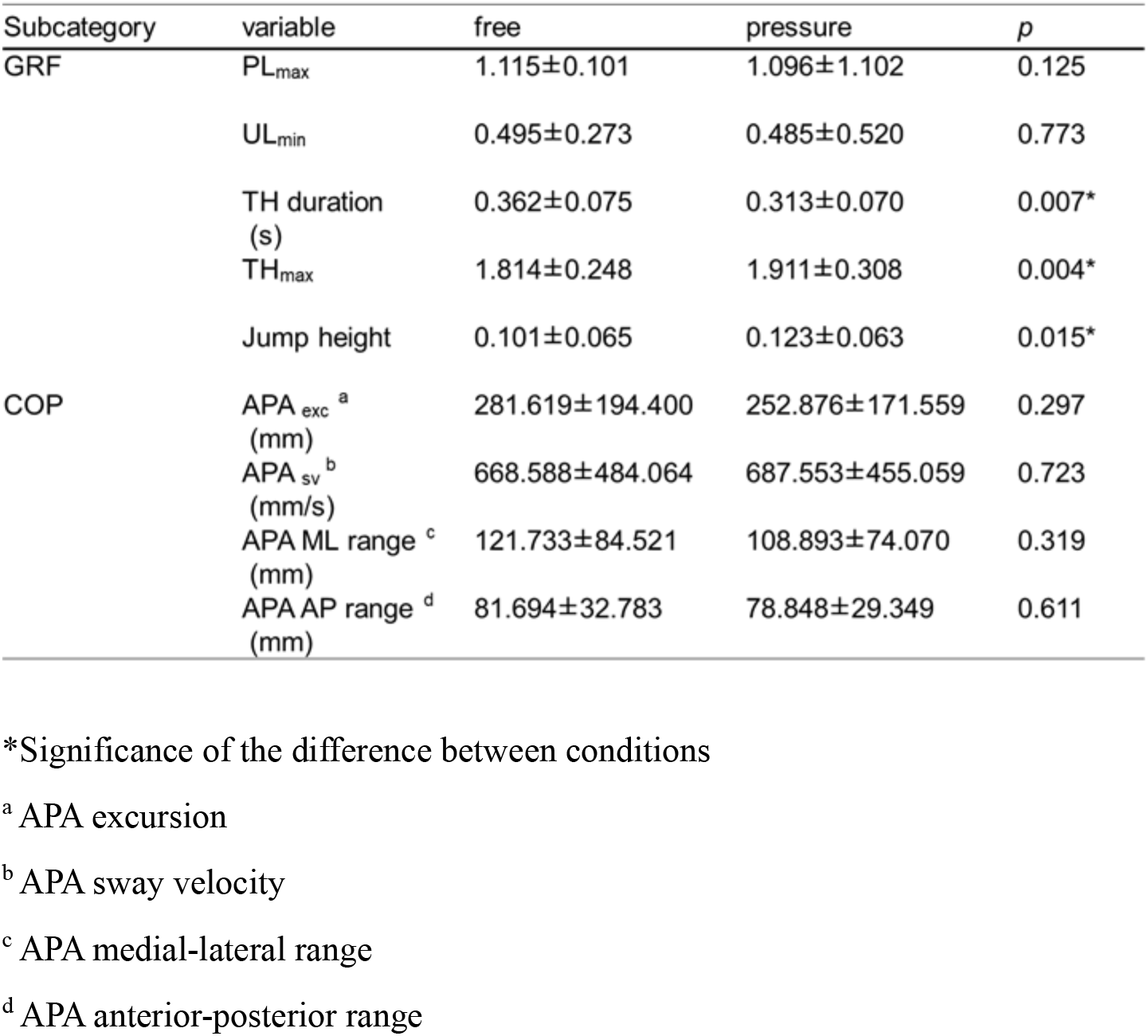
Means ± SDs of the Kinetics variables.

| Subcategory | variable | free | pressure | *p* |
| --- | --- | --- | --- | --- |
| GRF | PL<sub>max</sub> | 1.115±0.101 | 1.096±1.102 | 0.125 |
| | UL<sub>min</sub> | 0.495±0.273 | 0.485±0.520 | 0.773 |
| | TH duration (s) | 0.362±0.075 | 0.313±0.070 | 0.007* |
| | TH<sub>max</sub> | 1.814±0.248 | 1.911±0.308 | 0.004* |
| | Jump height | 0.101±0.065 | 0.123±0.063 | 0.015* |
| COP | APA<sub>exc</sub><sup>a</sup> (mm) | 281.619±194.400 | 252.876±171.559 | 0.297 |
| | APA<sub>sv</sub><sup>b</sup> (mm/s) | 668.588±484.064 | 687.553±455.059 | 0.723 |
| | APA ML range<sup>c</sup> (mm) | 121.733±84.521 | 108.893±74.070 | 0.319 |
| | APA AP range<sup>d</sup> (mm) | 81.694±32.783 | 78.848±29.349 | 0.611 |

\*Significance of the difference between conditions
<sup>a</sup> APA excursion
<sup>b</sup> APA sway velocity
<sup>c</sup> APA medial-lateral range
<sup>d</sup> APA anterior-posterior range

**Table 2.**
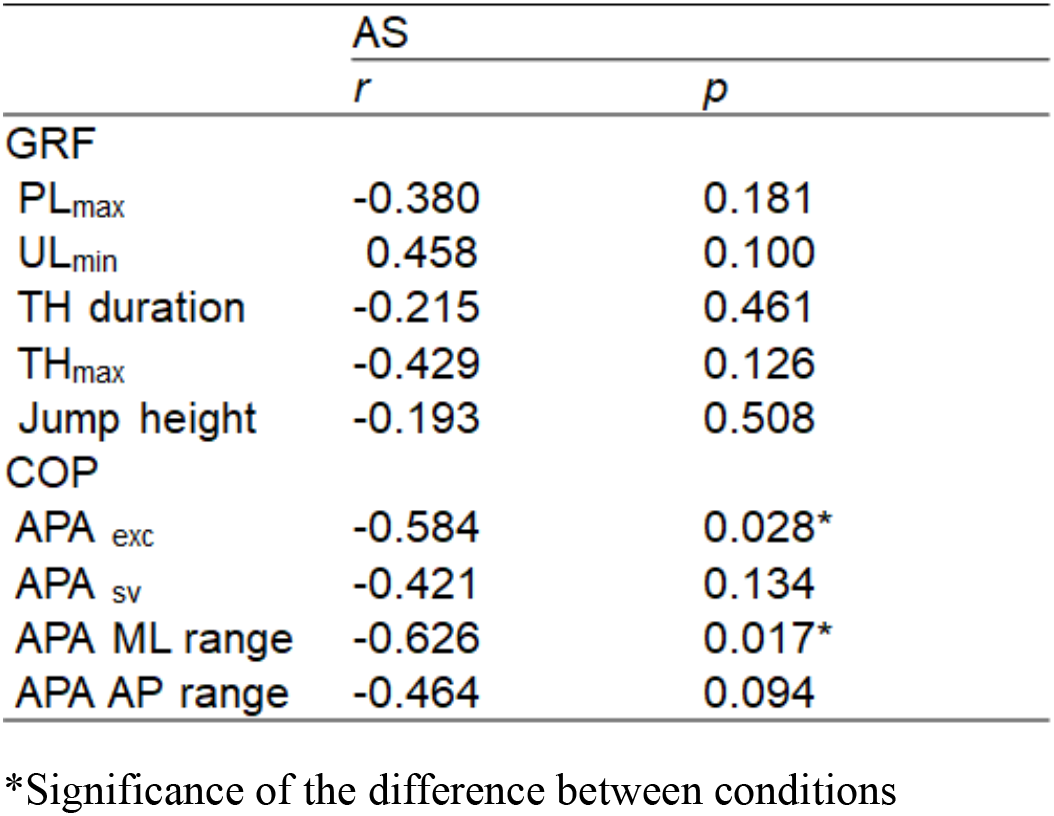
Pearson correlation coefficients between AS (f-p) and kinetics (f-p)

**Figure 2.**
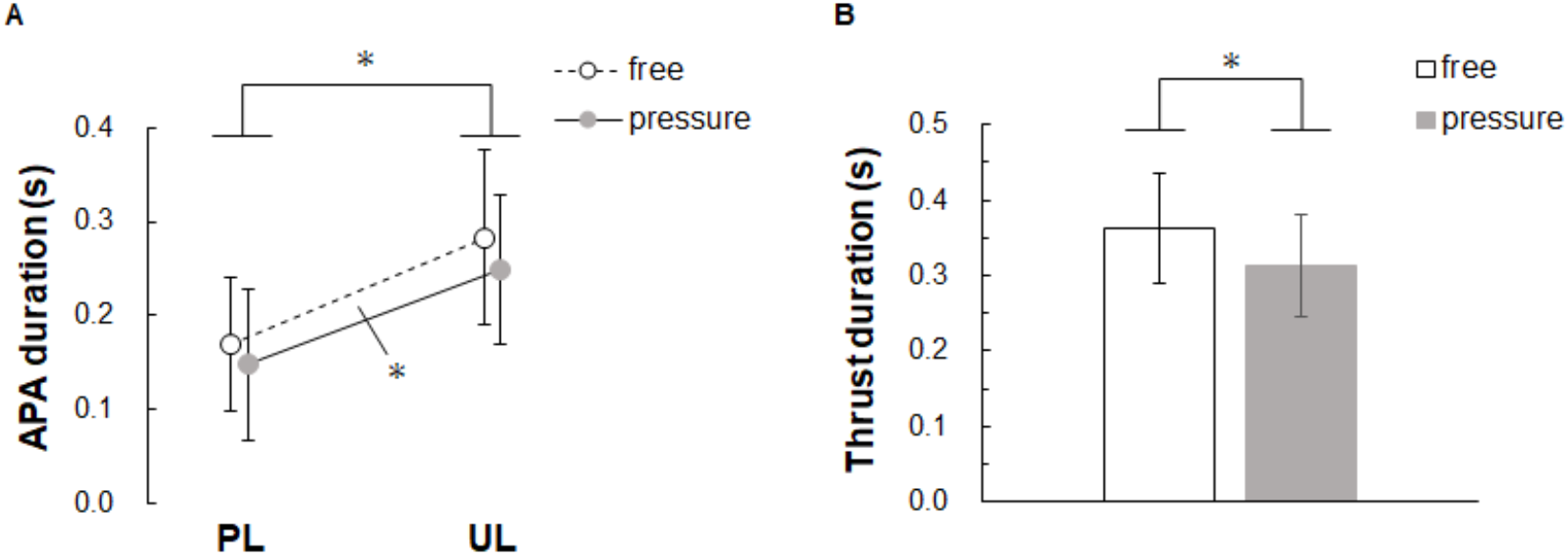
(A) the duration in the APA phase and (B) the duration in the TH phase are shown. *Error bars* represent between participant SD and *asterisks* denote a significant difference between phases or conditions (\**p* < .050).

For the thrust phase, The pairwise t-test revealed that the difference between conditions was significant for TH duration (t(13) = 3.1819, p = .007, Fig. 2B). Compared to the free condition, the pressure condition shortened the TH duration (free: 0.362±0.3786, pressure: 0.313±0.3052).

Similar to the TH_max_ and Jump height value, the pairwise t-test revealed that the difference between conditions was significant (t(13) = 2.7882, p = .015, t(13) = 3.5013, p = .004 respectively, Fig. 3A, B). Compared to the free condition, the pressure condition increased the TH_max_ and the Jump height value (free: 1.814±1.8100, pressure: 1.911±1.8644, free: 0.101±0.0895, pressure: 0.123±0.1201).

**Figure 3.**
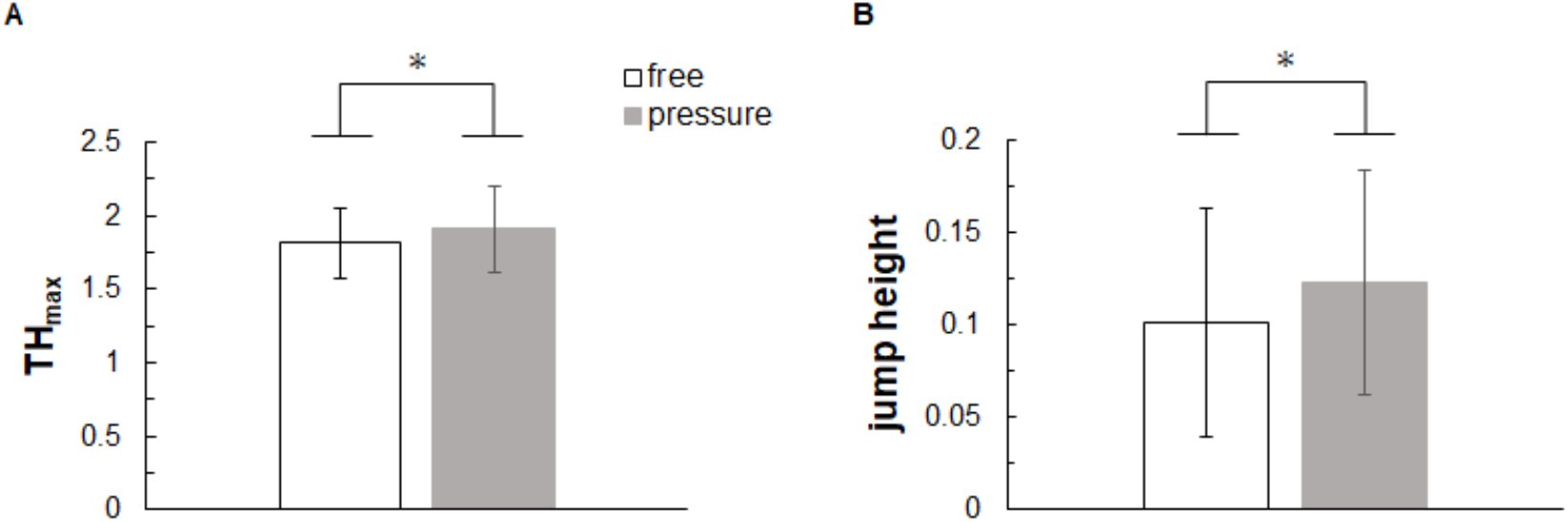
Kinetics variables obtained from jump shot movement. (A) maximum GRF during the TH phase and (B) jump height of jump shots are shown. *Error bars* represent between participant SD and *asterisks* denote a significant difference between conditions (\**p* < .050).

### 3.4. Relationship between AS and Kinetics variables

The correlation between the difference in AS between conditions (f-p) (ΔAS) and the difference in kinetics variance between conditions (f-p) are shown in Table 2. The ΔAS showed a significant negative correlation with the difference in APA excursion between conditions (ΔAPA excursion) (r = -0.584, p = .028, Fig. 4A). The greater the decrease in AS from free to pressure condition, the larger the decrease in the difference value of APA excursion between the two conditions.

**Figure 4.**
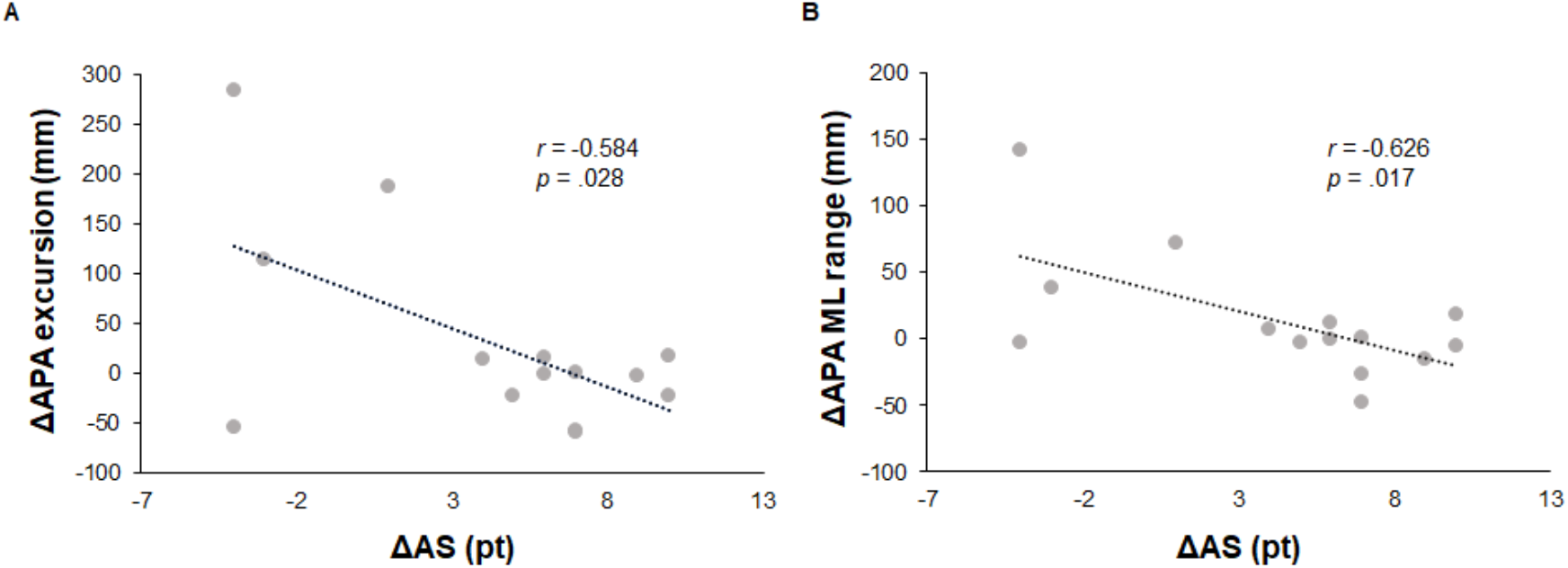
Relationship between the changes (f-p) in each variable across conditions. (A) ΔAS and ΔAPA excursion. (B) ΔAS and ΔAPA medial-lateral range (APA ML range).

Similar to the difference in APA ML range (ΔAPA ML range), there was a significant negative correlation with the ΔAS (r = -0.626, p = .017, Fig. 4B). The greater the decrease in AS from free to pressure condition, the larger the decrease in the difference value of APA ML range between the two conditions.

## 4. Discussion

The purpose of this study was to examine whether APAs are altered by pressure from a shot blocker and to investigate the relationship between APAs and jump shot performance in the free and pressure conditions.

In both the free and pressure conditions set in this study, the subjective pressure intensity of the participants was evaluated using the self-reported VAS score. The result of the subjective pressure intensity was significantly higher in the pressure condition than in the free condition. In addition, we found that AS considerably dropped in the pressure condition. Previous studies have shown that AS of jump shot decreases due to the presence of a shot blocker [16,17]. From these results, it is considered that the manipulation of pressure intensity using a shot blocker in the two conditions set in this study was successful.

The APA duration was shorter in the pressure condition compared to the free condition. This result supports the hypothesis that the APA duration will be shortened by the pressure from the shot blocker. For pressure from the shot blocker, the TH phase was shortened as the participants tried to perform the jump shot relatively quickly, which could be due to that the APA phase was shortened. In previous studies, voluntary movements such as jump shots are planned movements commanded by the feedforward model [22]. In other words, both are integrated at the planning stage before being output as motor commands. The previous study has demonstrated that trying to quickly perform only part of a pre-planned motor sequence affects the entire sequence [23]. Therefore, it is considered that the total duration of the jump shot motion is shortened along with the shortened APA phase.

The decrease in the AS under pressure from the shot blocker can be explained by Fitts law [24], which is a speed-accuracy trade-off whereby movement accuracy is increasingly compromised as motion speed goes up. In other words, it is considered that the AS would decrease as faster operation is required under the pressure condition. The following examines the factors that reduce accuracy and the APA duration due to the requirement for quick movements.

The APAs in the jump shot task in this study were considered to have two functions. One function is to generate reaction forces. Previous studies have demonstrated that SSC movements tend to shorten the time between stretch and shortening phases to enable a faster operation [25]. In addition, Komi (1984) has shown that shortening the time between the stretch and shortening phases increases muscle power output [30]. In this study, it is considered that the shorter APA and TH durations may increase the TH_max_ and Jump height values. Other previous studies have demonstrated that changes in muscle power affect precision and fine control of motions [26,27]. Therefore, excessive leg muscle output may cause a decrease in jump shot performance.

The other function of APAs is postural stability. Similar previous studies have indicated the association between balance ability and shooting accuracy [9–11]. When an operation is performed quickly, the COP displacement decreases in the APA phase as the APA duration is shortened, but the COP displacement increases during the main motion [19]. That is, it has been shown that the shorter APA duration destabilizes posture in the subsequent main motion. Likewise, under the pressure condition in this study, we presumed that COP variables would change due to the pressure from the shot blocker. However, no difference between conditions was observed regarding COP variables in this study. These results did not support our hypothesis.

A previous study has shown that APAs are established on previous experience and can be acquired through learning [28]. Therefore, COP control depends on both the level of individual motor skills and experience according to motor learning. In this study, there was an 11 point difference between participants regarding the AS under the pressure condition. Such differences in skill levels might make it difficult to detect the statistically significant difference. This seems to be substantiated by the findings that the AS negatively correlated with APA excursion and APA ML range. That is, those whose postural stability hardly changed despite the shot blocker’s pressure showed less decrease in the AS. A previous study has shown that sport-specific training causes adaptation of postural control [29]. That is, highly skilled athletes may improve postural control through long-term training and be capable of suppressing body sway. Therefore, those capable of maintaining postural stability even within the limited time under the pressure condition, as in this study are considered less likely to decline the AS.

This study focused on the jump included in the jump shot motion and detected APAs from the ground reaction force. However, further investigation is needed to ascertain changes in kinematics variables. This study is the first finding to further our understanding of postural mechanisms in practical settings. Future studies are necessary to investigate the effects of anticipatory postural adjustments on jump shot performance for participants of various skill levels for further application to practical settings.

## Conclusion

This study revealed that APA duration in the jump shot motion was shortened under pressure from the shot blocker. Incorporating jump shot practice under pressure from a shot blocker may help players adapt to shortened APA durations. Additionally, measuring each player’s APA duration and associated changes and analyzing their impact on performance could serve as indicators for improvements.

### Fund

This research was supported by JSPS KAKENHI Grant number 22H03492 and 23K10757.

## Notes

### Competing Interest Statement

The authors have declared no competing interest.

